# Embryonic exposure to valproic acid and neonicotinoid deteriorates the developmental GABA switch and impairs long-term potentiation in the local circuit of intermediate medial mesopallium of chick telencephalon

**DOI:** 10.1101/2024.09.15.613159

**Authors:** Toshiya Matsushima, Noriyuki Toji, Kazuhiro Wada, Hiroki Shikanai, Takeshi Izumi

## Abstract

Embryonic exposure to valproic acid (VPA) and imidacloprid (IMI, a neonicotinoid insecticide) impairs filial imprinting in hatchlings, and the deteriorating effects of VPA are mitigated by post-hatch injection of bumetanide, a blocker of the chloride intruder NKCC1. Here, we report that these exposures depolarized the reversal potential of local GABAergic transmission in the neurons of the intermediate medial mesopallium (IMM), the pallial region critical for imprinting. Furthermore, exposure increased field excitatory post-synaptic potentials in pre-tetanus recordings (fEPSPs) and impaired long-term potentiation by low-frequency tetanic stimulation (LTP). Bath-applied bumetanide rescued the impaired LTP in the VPA slices, whereas VU0463271, a blocker of the chloride extruder KCC2, suppressed LTP in the control slices, suggesting that hyperpolarizing GABA action is necessary for the potentiation of excitatory synaptic transmission. However, the transcriptional profiles of IMM slices did not support the expected increase in the NKCC1/KCC2 ratio, suggesting a potential modification of post-transcriptional processes. Instead, exposure to both VPA and IMI downregulated several transcriptional regulators (FOS, NR4A1, and NR4A2) and upregulated the RNA component of signal recognition particles (RN7SL1). As a limited set of response genes were shared, VPA and IMI could cause common neuronal malfunctions via distinct molecular cascades.

## Introduction

Chemicals such as valproic acid (VPA) and neonicotinoid insecticides have been studied as potential environmental risk factors for developmental disorders, including autism spectrum disorder (ASD) when exposed during pregnancy (Moore et al. 2000, Rasalam et al. 2005, Bromley et al. 2013, Christensen et al. 2013 for VPA; Keil et al. 2014; Gunier et al. 2017; von Ehrenstein et al. 2019; Ongono et al. 2020 for neonicotinoids). Accordingly, in model animals, embryonic exposure to these agents impairs post-natal development of social behaviors in rodents (Meador and Loring 2013, Mabunga et al. 2015, Ergaz et al. 2016, Sano et al. 2016, Nicolini and Fahnestock 2018, Chaliha et al. 2020) and domestic chicks (Nishigori et al. 2013, Sgadò et al. 2018, Lorenzi et al. 2019, Adiletta et al. 2021, Matsushima et al. 2022). VPA acts as an inhibitor of histone deacetylase (Phiel et al. 2001), whereas neonicotinoids such as imidacloprid (IMI) are potent agonists of the nicotinic acetylcholine receptor (nAChR) (Costas-Ferreira and Faro 2021). How do these agents with different chemical actions cause similar behavioral maldevelopment?

Despite diverse and heterogeneous genetic and environmental risk factors, ASD phenotypes can arise by an imbalance between excitatory and inhibitory synaptic transmission (E-I imbalance) in cortical and subcortical networks for processing perception, memory, and emotions (Rubenstein and Merzenich 2003, Nelson and Valakh 2015, Gilbert and Man 2017). Enhancement of the hyperpolarizing GABA action by bumetanide (a blocker of NKCC1, Na-K-Cl cotransporter 1 as chloride intruder) appeared to produce positive outcomes in some cases of ASD (Hadjikhani et al. 2018, Fernell et al. 2021, Wang et al. 2021, Delpire and Ben-Ari 2022). Note however that recent phase-2 trials failed to support this efficiency (Sprengers et al. 2021, van Andel et al., 2023). The possible efficacy of bumetanide treatment has also been suggested in a mouse model of fetal alcohol spectrum disorders (FASD; Skorput et al. 2019). E-I imbalance can produce behavioral impairments also in precocial birds. Specifically, when applied to newly hatched chicks, bumetanide rescued imprinting deterioration caused by embryonic exposure to VPA (Matsushima et al. 2022).

Newly hatched chicks could be a valid animal model to study ASD, because they are innately predisposed to pay visual attention to animate objects and face-like configuration (Vallortigara et al. 2005, Miura and Matsushima 2016, Miura et al. 2012, Miura et al. 2020 for biological motion; Rosa-Salva et al. 2010, 2011, 2019 for face-like configuration) as of human newborns (Johnson 1992, Johnson 2005, Simion et al. 2008, Pavlova 2012, Pavlova et al. 2017); for recent reviews, see Matsushima et al. 2024, Sgadò et al. 2024, Kobylkov and Vallortigara 2024. In addition to the surface validity, chicks have constructive validity as an ASD model. Successful filial imprinting is associated with an enhanced inflow of thyroid hormones (triiodothyronine, T_3_) into the dorsal pallial network, which elongates the sensitive period of imprinting (Yamaguchi et al. 2012). Furthermore, the higher the T_3_ level, the more disposed the chicks are to social objects bearing vivid motion animacy (Takemura et al. 2018, Miura et al. 2018, Lorenzi et al. 2021). Finally, bath-applied T_3_ enhances GABAergic transmission in *in vitro* slice preparations of the hatchling’s telencephalon (Saheki et al. 2022), suggesting a critical role of the E-I balance for social attachment formation.

Given the critical importance of balanced GABA transmission during the early stages of development, how does this appear to be maladaptive throughout postnatal life? Compensatory mechanisms have been assumed to contribute to the adaptive reorganization of neural circuits (Nelson and Valakh 2015), yet it remains unknown why these mechanisms fail in ASD cases. To address this issue, we examined how embryonic exposure to these neurodevelopmental toxicants alters synaptic function in the intermediate medial mesopallium (IMM), which is the brain region responsible for imprinting (Horn 1985, Horn 1998, Horn 2004). In addition to the biochemical, morphological, and neurophysiological changes in the IMM associated with imprinting (Horn et al., 1985, McCabe and Horn 1994, Horn et al. 2001, Solomonia et al. 2003), local excitatory synaptic transmission shows long-term potentiation (LTP) by low-frequency tetanic stimulation (Bradley et al. 1991, Bradley et al. 1999). Notably, LTP is suppressed when GABAergic transmission is blocked by bicuculline (Yanagihara et al. 1998), suggesting that plastic changes in excitatory synapses require GABA action. However, when the post-synaptic neuron is hyperpolarized, the same tetanic stimulation instead induces a long-term depression (Matsushima and Aoki 1995), indicating bidirectional Hebbian nature of the plastic changes. Here, we report a strong association among the GABA switch, synaptic E-I balance, and LTP in the emergence of ASD-like phenotypes following embryonic exposure to environmental toxicants in domestic chicks. In particular, we examined the effects of blockers of chloride cotransporters (bumetanide for NKCC1 and VU0643271 for KCC2, K-Cl cotransporter 2 as chloride extruder) on LTP. Although the subcellular mechanisms are yet to be elucidated, the hyperpolarizing GABA switch could be critical for the developmental control of the E-I balance through homeostatic synaptic plasticity.

## Results

### Effects on the developmental GABA switch

Whole-cell patch recordings were successfully obtained in 164 neurons from 41 chicks (post-hatch 1 day, P1; 12-36 hours post-hatch; 15 males and 26 females) and 8 embryos (embryonic 18 and 20 days, E18 and E20; unsexed); 79 neurons by single-electrode discontinuous voltage clamp (SEVC) and 85 neurons by conventional voltage clamp (VC). As chicks usually hatch on E21 = P0, peri-hatch recordings were made at two-day intervals. In 143 of the 164 neurons, current responses showed a clear dependence on the holding membrane potential (V_holding_) around the resting membrane potential (V_rest_), and the reversal potential (V_rev_) was reliably determined at a precision of 1-3 mV by systematically changing V_holding_. In the other 21 neurons, V_rev_ was not measurable either because the neuron was directly excited to fire by a low-intensity stimulation subthreshold for post-synaptic responses, or because the response was not dependent on V_holding_.

The traces in **Fig. 1B** show examples recorded from a control P1 neuron. In normal Krebs solution (**Fig. 1Ba**), the synaptic currents were composed of an initial inward current followed by a net outward current when V_holding_ was set above V_rest_. The outward component (presumptive GABA component; the onset indicated by vertical green lines) was reversed to a net inward component at hyperpolarized V_holding_. The blockade of fast excitatory synaptic transmission by DNQX in the bath (**Fig. 1Bb**) unmasked the GABAergic late component with clear V_holding_ dependence. The reversal potential (V_rev_ = -65 mV) was determined as the V_holding_ where the earliest component of the GABA current was nullified (I_ipsc_ = 0 pA; horizontal red lines). Comparisons among 3 groups of unexposed control (E18, E20, and P1) revealed a hyperpolarizing developmental shift in V_rev_ (**Fig. 1Ca**), with slight changes in V_rest_ (**Fig. 1Cb**); see Supplementary **Fig. S1** for the V_rev_ vs. V_rest_ plots and statistical examinations using a linear fitting analysis. As shown below (**Fig. 4Ba**), qPCR analysis confirmed the developmental decline in the ratio of slc12a2 to slc12a5, the genes that encode NKCC1 and KCC2 respectively.

**Fig. 1.**
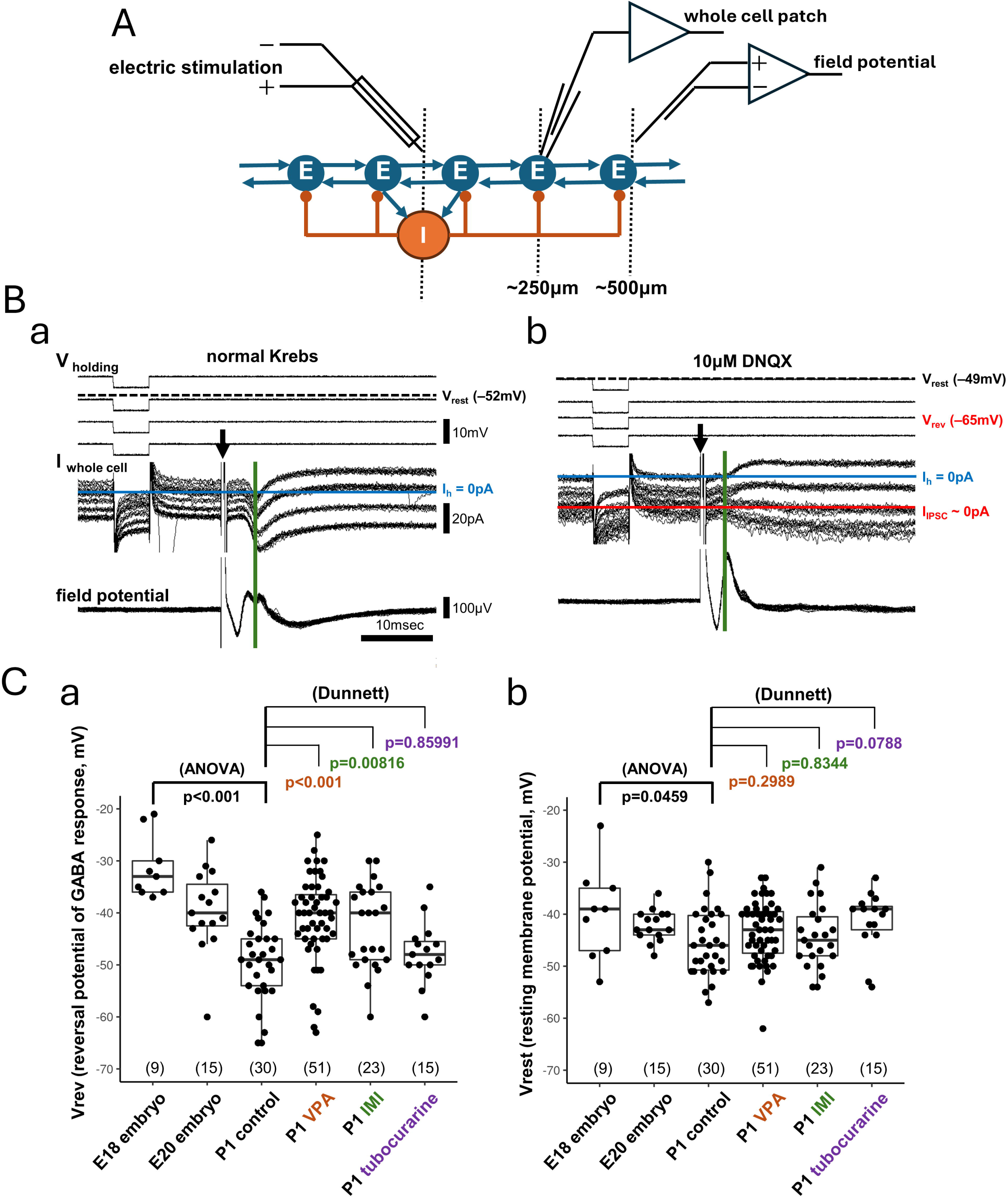
Whole-cell voltage recording from intermediate medial mesopallium (IMM) neurons revealed the reversal potential of local GABAergic fast transmission. **A)** Schematic diagram of the estimated local network of IMM composed of excitatory and inhibitory neurons (E and I, respectively). **B)** Representative recording of whole-cell current (I _whole cell_) by voltage-clamp mode in normal Krebs **(a)** and after DNQX was added to the bath perfusate **(b)**; superimposed traces (>10 for each level of the holding potential V_holding_). Stimulus intensity: 111 μA (**a**) and 200 μA (**b**); duration: 400 μsec; intervals: 2 sec. Vrev (reversal potential) was determined at the level of null initial synaptic current (I_IPSC_ = 0, red line). Vrest (resting potential) was determined at the level of null holding current (I_h_ = 0, blue line). **C)** Comparisons of Vrev **(a)** and Vrest **(b)** among groups. One-way ANOVA was applied to the three control groups (E18, E20, and P1); a rapid hyperpolarizing shift occurred during four days of the pre-/neo-natal period. Multiple comparisons were made using Dunnett’s method to the P1 groups (VPA [7 and 35μmole per egg], IMI [50 and 250μg/egg], and tubocurarine [0.2mg/egg] against control as the reference); significantly depolarized Vrev was found in VPA and IMI groups, but not in tubocurarine. Numbers in parentheses indicate the number of recorded neurons. See **Fig. S1** for the dose response.

The V_rev_ effects paralleled the behavioral effects on imprinting reported previously (Matsushima et al. 2022), namely, impairment by VPA and IMI but not by tubocurarine. V_rev_ was depolarized when chicks were exposed to VPA and IMI on embryonic day 14 (E14) but not tubocurarine (**Fig. 1Ca**). We examined the effects of VPA and IMI treatments in two doses (7 and 35 μmole for VPA, and 50 and 250 μg for IMI per egg); data obtained from these two dose levels were merged because of considerable overlaps (Supplementary **Fig. S1**). Multiple comparisons using Dunnett’s method revealed differences in VPA and IMI compared to the P1 control group, whereas no significant effects were found on V_rest_ (**Fig. 1Cb**).

Bath applied bumetanide acutely hyperpolarized V_rev_ without changes in V_rest_ in VPA slices; **Fig. 2A** shows an example. With DNQX in the bath, V_rev_ was around V_rest_ (-43mV). When bumetanide (blocker of NKCC1; 20 μg/mL or 55 μM) was added to the DNQX perfusate, V_rev_ was hyperpolarized by 10 mV in 15-20 min (**Fig. 2Ab**). The current component was GABAergic as it was blocked by bicuculline (**Fig. 2Ac**). **Fig. 2C** shows the population data. Changes in V_rev_ and V_rest_ (ΔV_rev_ and ΔV_rest_) recorded at 15-25 min after the onset of bumetanide perfusion are plotted and tested by one-sample t-test against the null hypothesis of ΔV = 0 mV. Note that vehicle of the bumetanide solution (pH = ca. 8.2 due to addition of NaOH) had no significant effects. Therefore, IMM neurons are candidate targets of bumetanide for the acute rescue of imprinting in the VPA exposed chicks (Matsushima et al. 2022). On the other hand, VU0463271 (blocker of KCC2; 1 μM) hyperpolarized V_rev_ in control slices, again without significant changes in V_rest_; **Fig. 2B** shows example traces, and **Fig. 2C** the population data. Therefore, we examined the effects of bumetanide and VU0463271, applied under the same conditions (concentration and duration of treatment), on synaptic plasticity measured by field potential recording.

**Fig. 2.**
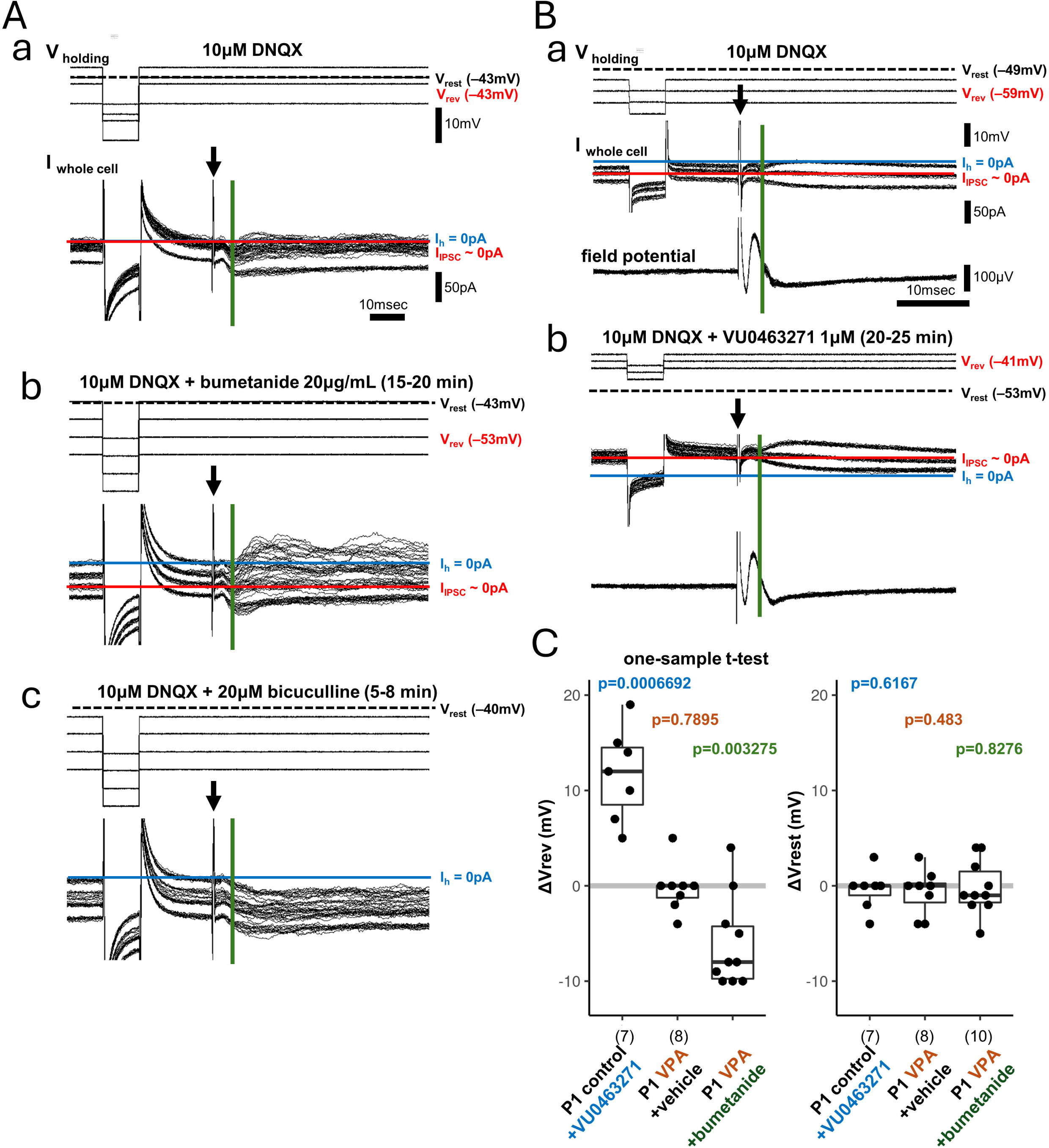
Acute effects of bumetanide (NKCC1 blocker) and VU0463271 (KCC2 blocker) on Vrev and Vrest of VPA-treated and control chicks, respectively. Representative recordings (**A** and **B**) and population data (**C**). **A**) In IMM slice obtained from a VPA-treated chick, Vrev was determined in DNQX-Krebs (**Aa**), and in bumetanide added to the perfusion for 15-20 min (**Ab**); hyperpolarization shift occurred at ΔVrev = -10mV. Replacement of bumetanide with bicuculine completely vanished the voltage-sensitive current, leaving a lasting inward current (**Ac**). **B)** In the control slice, the addition of VU0463271 to the perfusate Krebs caused a depolarizing shift of Vrev. **C)** ΔVrev and accompanying ΔVrest are shown in each group; a one-sample t-test was repeated for each.

### Effects on the synaptic E-I balance and LTP

Extracellular field potential recordings were successfully obtained from 70 slice preparations from 32 chicks (P1: 14 males and 18 males). Presynaptic fiber volleys were reliably recorded in 68 slices; in the other two slices (exposed to IMI), large stimulus artifacts obscured the pre-synaptic volleys.

**Fig. 3B** illustrates how presynaptic volleys and fEPSP were measured. The stimulating and recording electrodes were placed such that a biphasic compound spike (presynaptic fiber volley) was followed by a small positive deflection (shoulder) on the decaying slope (pre-tetanus phase, 60 superimposed traces for 10 min). Low-frequency tetanic stimulation (5 Hz for 60 s, 300 pulses) was applied to induce LTP, as reported previously (Yanagihara et al. 1998). Post-tetanus recordings lasted for 30 min at 10 s intervals; the superimposed traces obtained in the last 10 min (60 traces) of the post-tetanus phase are shown. The slice was subsequently perfused with the DNQX Krebs to reveal the onset (blue dashed vertical lines) and amplitude of the monosynaptic component of EPSP (red dashed vertical lines), and the base of the DNQX sensitive component (DNQX base; dashed horizontal line) was determined; average traces were enlarged and superimposed on the right. In the slice shown here, the fEPSP onset occurred 2.3 msec after the stimulus, and its amplitude was measured at 3.1 msec. The fEPSP was normalized to the pre-synaptic volley recorded during the pre-tetanus phase. The fEPSP onset ranged from 1.3 to 3.1 msec (2.26 ± 0.04 msec, mean ± s.e.m., n=70), and its amplitude was measured at the latency ranging from 0.3 to 1.7 msec (0.97 ± 0.03 msec, mean ± s.e.m., n=70).

**Fig. 3.**
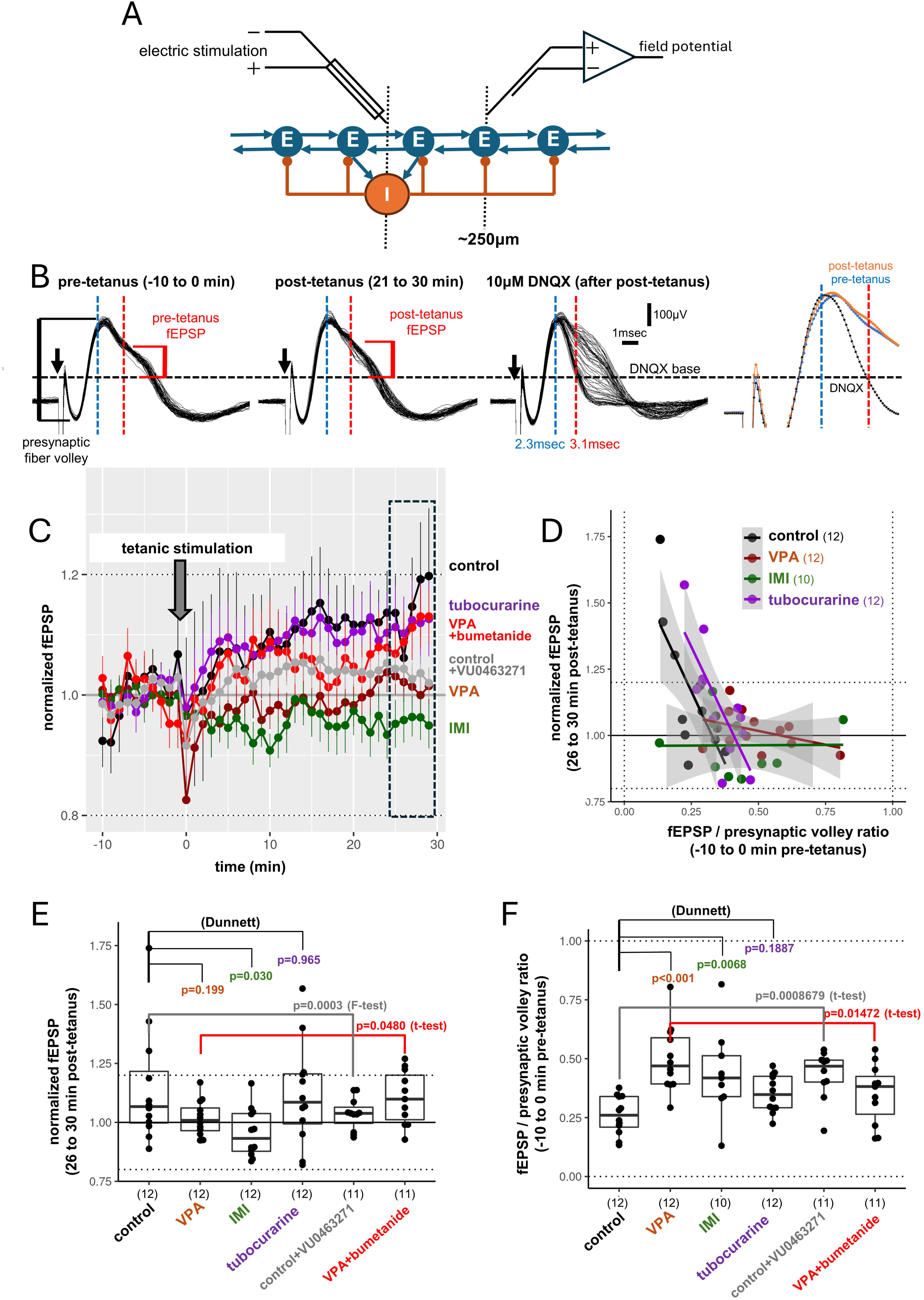
Potentiation of excitatory post-synaptic responses was examined in IMM slices. **A)** Schematic diagram. **B)** Representative recording of field EPSPs (fEPSPs); superimposed pre-tetanic traces (left, 60 traces for 10 min), post-tetanic traces (middle, 60 traces for 10 min), and traces recorded in DNQX-Krebs (right, 36 traces for 6 min). Pre-synaptic fiber volleys were measured as the amplitude of the initial bi-phasic component just after the local electrical stimulation. The amplitude of the fEPSPs was measured at 0.8 msec (red dashed lines) from the onset point of the DNQX-sensitive component (blue dashed lines). The traces recorded at 5-6 min after the DNQX application yielded the base level (DNQX base, black dashed horizontal line). **C)** Time course of the fEPSPs before and after the low-frequency tetanic stimulation (5Hz x 300 pulses, downward arrow) after normalization by the pre-tetanic level. Dots and vertical lines indicate the mean and s.e.m. (6 traces x 12 or 11 experiments per min). **D)** Normalized fEPSP changes induced by tetanic stimulation (LTP, y-axis) was plotted against the fEPSP amplitude recorded pre-tetanus (x-axis) in four groups of slices. See Supplementary **Fig. S2** for the acute effects of bumetanide and VU0463271. **E)** Normalized fEPSPs were compared during the last 5 min (26-30 min post-tetanus, dashed box). Multiple comparison using Dunnett’s method was applied to VPA [35 μmole per egg], IMI [250 μg/egg], and tubocurarine [0.2 mg/egg] against control data; Multiple comparisons revealed significant differences only in IMI. Repeated pairwise comparisons revealed significant differences between control vs. control+VU0463271 (significant by F-test) and between VPA vs VPA+bumetanide (t-test). **F)** The ratio of the fEPSP amplitude to the pre-synaptic fiber volley was compared among groups. Significant difference was found in VPA and IMI, but not in tubocurarine. Pairwise comparisons revealed significant differences between control vs. control+VU0463271 and between VPA vs. VPA+bumetanide.

The fEPSP amplitude was normalized by the pre-tetanus average and plotted against the time starting from -10 min (at the point after the recording was stabilized) until 30 min post-tetanus (**Fig. 3C**). The mean of six consecutive traces (at 10 s intervals, i.e., 1 min) was further averaged over the number of slice preparations (12 or 11 slices) for each time point, with s.e.m. of 72 or 66 traces as the vertical error bars. Note that intergroup differences emerged as early as 5-10 min post-tetanus and monotonically increased afterwards. If the last 5 min period (26-30 min post-tetanus) is focused, control and tubocurarine slices showed comparable LTP, whereas the slices exposed to VPA (35 μmole per egg) and IMI (250 μg per egg) failed to show LTP. Furthermore, LTP was resumed if the VPA slices were treated by bumetanide in the bath (55 μM) starting at ca. 20 min before the pre-tetanus recordings (VPA+bumetanide). On the other hand, pre-treatment of control slices with VU0463271 (1 μM) for ca. 20 min suppressed LTP (control+VU0463271).

The normalized amplitude of post-tetanus fEPSP during the last 5 min was averaged for each slice and compared among the groups (**Fig. 3E**). VPA and IMI slices, but not tubocurarine, failed to show LTP at the population level, although a statistically significant level appeared only in IMI according to the ANOVA test of the Dunnett’s method. Direct pairwise comparisons between control vs. control+VU0463271 and VPA vs. VPA+bumetanide yielded significant differences using the F-test and t-test, respectively. Therefore, we conclude that LTP correlated with the developmental GABA switch (**Fig. 1Ca**) at the group population level.

The fEPSP amplitude normalized by the pre-synaptic fiber volley (recorded in the pre-tetanus phase) showed a nearly negative image of LTP, where the VPA and IMI groups were higher than the control and tubocurarine groups (**Fig. 3F**). In addition, at the level of individual slices (**Fig. 3D**), a higher LTP appeared when the fEPSP/pre-synaptic volley ratio was lower, and the interaction with the group was significant. Two-way ANOVA revealed a significant main effect of fEPSP (F=10.6436, Df=1, p=0.0002) and interaction with the treatment (F=9.0488, Df=3, p=0.0001) but the main effect of the treatment was not significant (F=0.14954, Df=3, p=0.2312); for detailed statistics see Supplementary materials. The VPA and IMI slices, but not the tubocurarine slices, showed a statistically higher fEPSP/pre-synaptic volley ratio after ANOVA using the Dunnett’s method (**Fig. 3F**). Direct pairwise comparisons between the control and control+VU0463271, as well as VPA vs. VPA+bumetanide, yielded significant differences by t-test. Therefore, we conclude that embryonic exposure to VPA and IMI increases DNQX-sensitive excitatory synaptic responses in a manner coupled with LTP impairment.

### Effects on the gene expression profiles, RNAseq, and qPCR analyses

To search for the underlying molecular events of the deterioration in the GABA-switch and impaired LTP, gene expression profiles were examined by RNA-seq using the punched tissue of the IMM studied for whole-cell recording. Slice preparations obtained from six chicks (all females, two chicks each for the control, VPA, and IMI groups) were chosen such that group-typical V_rest_ and V_rev_ values were recorded (see the inlet table in **Fig. 4A**). After whole-cell recordings were made, 1 mm-sized punched tissues were sampled around the recording sites from four consecutive slices and mixed to represent each chick. Histological reconstruction revealed that the sampling was confined mostly within the IMM (**Fig. 4A**).

**Fig. 4.**
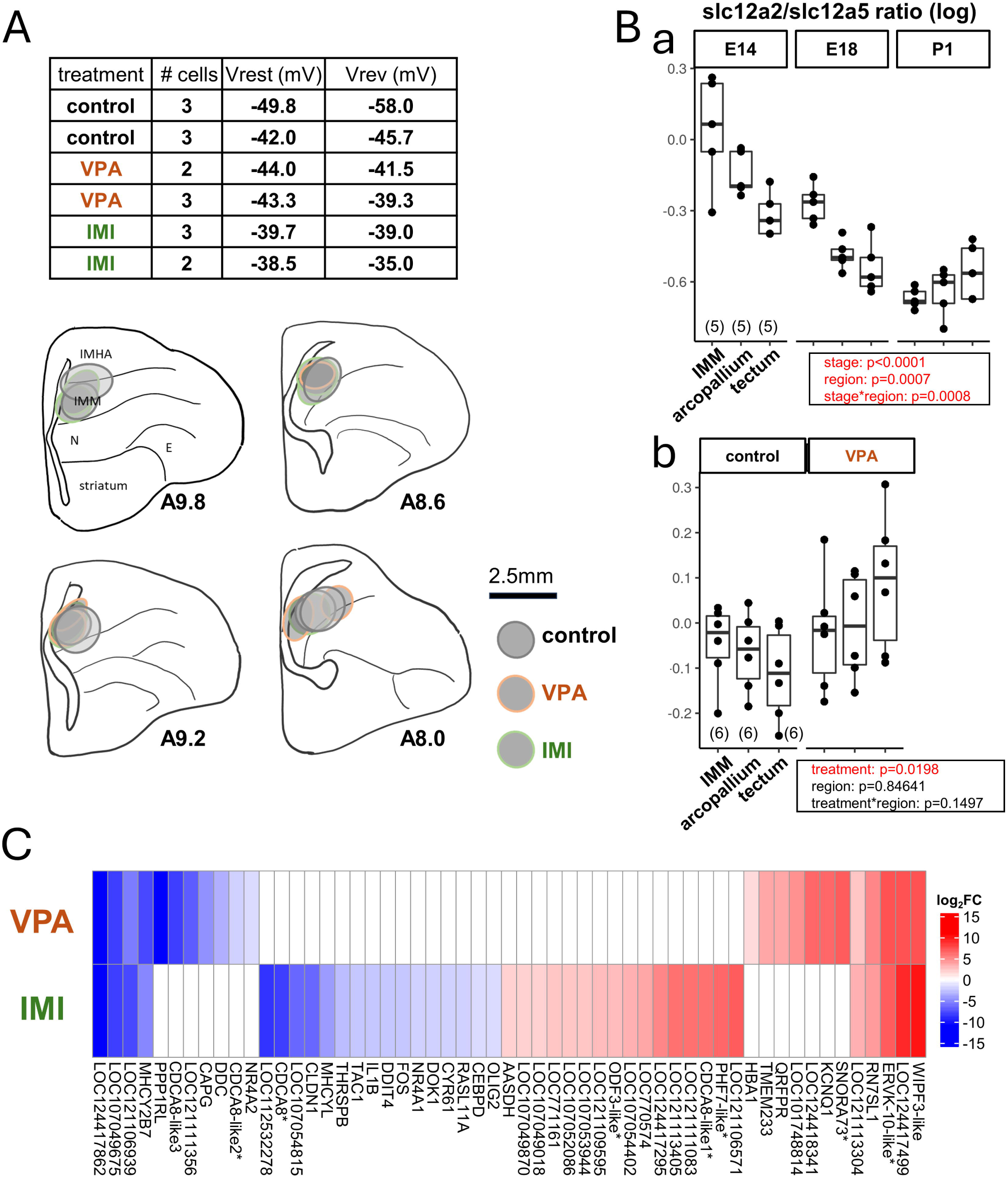
Gene expression profiles. **A)** Histological reconstruction of sampled regions for RNA-seq analysis in IMM. The inlet table shows averaged V_rest_ and V_rev_ recorded from 6 chicks: 2 controls, 2 VPA [35 μmole per egg], and 2 IMI [250 μg/egg]. Four pieces of punched tissue were sampled from each chick at A9.6 to A8.0; the AP level follows the chick brain atlas by Kuenzel & Masson (1988). **B)** Ratio of slc12a2 (NKCC1) for slc12a5 (KCC2) gene expression revealed by qPCR analysis in 3 brain regions, IMM, arcopallium, and optic tectum. Developmental changes from embryonic E14, E18, and post-hatch P1 (**Ba**) and the effects of E14 exposure to VPA were analyzed on P1 (**Bb**). Inlet boxes show the statistics. **C)** Expression fold changes in the genes with significantly different expression by FDR (fault discovery rate) < 0.1 for VPA and IMI. Upregulated and downregulated levels, compared to the control, are color coded in red and blue.

Statistically different gene expressions with a false discovery rate (FDR) below 0.10 are summarized in **Fig. 4C** for VPA and IMI; full lists are shown in Supplementary **Table S2**. Out of 25,467 genes (and 85,704 isoforms), small sets of 24 (for VPA) and 42 genes (for IMI) showed increased or decreased expression, respectively, compared with the control. Nine genes were overlapping between VPA and IMI. Two of these nine genes were annotated, namely upregulated RN7SL1 (RNA component of signal recognition particle 7SL1) and downregulated MHCY2B7 (major histocompatibility complex Y, class II beta 7); the rest were classified as LOC genes. Among the VPA and IMI sets, NR4A2 (VPA, down-regulated; nuclear receptor subfamily 4 group A member 2), NR4A1, and FOS (IMI, down-regulated) require particular attention, because de novo variants of NR4A2 are associated with several neurodevelopmental cases (Lévy et al. 2018, Singh et al. 2020, Song et al. 2022) and NR4A1 is associated with dendritic spine formation in cortical neurons (Chen et al. 2014, Li et al. 2016). However, NKCC1 and KCC2 were not included in the list of low-FDR genes.

Parallel to the developmental changes in V_rev_ (**Fig. 1Ca**), the expression ratio of the genes encoding the chloride cotransporters (ratio of slc12a2 for NKCC1 to slc12a5 for KCC2, on a log scale, **Fig. 4Ba**; qPCR analysis) showed a significant decline from E14 to E18 to P1 (Supplementary **Fig. S3** for further details). Punched tissue samples were prepared from three brain regions (IMM, arcopallium, and optic tectum) without electrophysiological experiments from five chicks at each stage, and gene expression profiles were analyzed using qPCR. Two-way ANOVA revealed a significant effect of development (p<0.0001) and brain region (p=0.0007), with a significant interaction (p=0.0008; see Supplementary for statistics).

Contrary to our expectations, the slc12a2/slc12a5 ratio did not significantly differ between the control and VPA groups in the qPCR dataset (**Fig. 4Bb**). Three brain regions were sampled, each from six control and six VPA chicks. A significantly higher ratio occurred in the VPA group compared to the control group (p=0.0198) without effects of region (p=0.84641) and interaction (p=0.1497). Note however the considerable overlaps of the ratio in IMM between VPA and the control. The RNA-seq dataset also showed non-significant differences in SLC12A2, SLC12A5, and their isoforms among the control, VPA, and IMI chicks (Supplementary **Table S3**). Despite their distinct effects on V_rev_ (**Fig. 1**), embryonic exposure to VPA and IMI did not alter the mRNA expression of NKCC1 or KCC2 in the IMM of hatchlings. This conclusion needs a careful consideration, because NKCC1 is expressed in many cell types, whereas KCC2 is neuron-specific (Kaila et al. 2014, Watanabe and Fukuda 2015, Löscher and Kaila 2022). Future studies must be conducted at the single-cell level.

## Discussion

The present results revealed the cellular events associated with the perinatal development of social behavior and its disorders by environmental toxicants; **Fig. 5** schematically illustrates the findings. Embryonic exposure to VPA and IMI alters developmental landscape (Matsushima et al. 2024), causing distinct gene expression patterns in the IMM of hatchlings that are critical for imprinting. The characteristic set of events accompanied at the neuronal level, namely, depolarized V_rev_, increased EPSP, and impaired LTP induction. The deteriorated GABA switch reduces the efficacy of the inhibitory action of GABA, thereby boosting the E-I imbalance. These events are not supposed to be linked to the impaired biological motion preference caused by the IMI exposure (Matsushima et al. 2022). Probably, impairments in the other sub-pallial visual pathways are responsible.

**Fig. 5.**
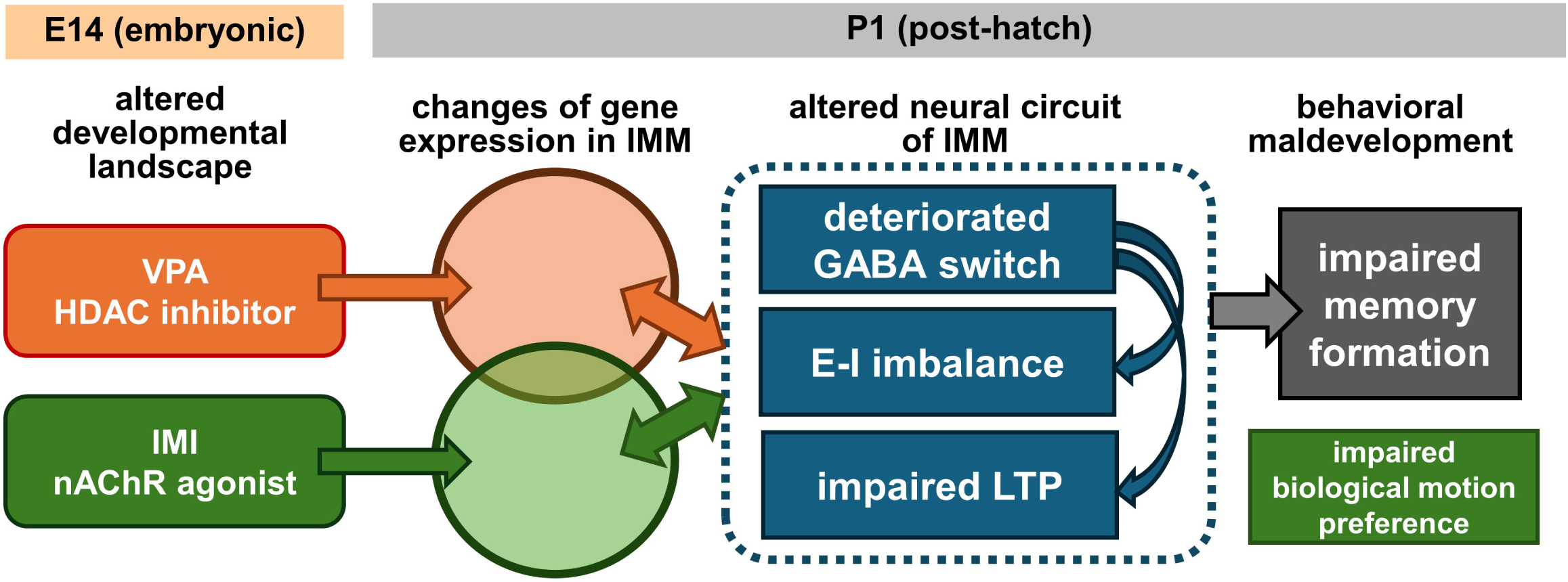
Schematic illustration of the cellular cascades from embryonic exposure to VPA and IMI, through gene expression changes and altered neural circuit of IMM, to impaired memory formation of imprinting.

To the best of our knowledge, the present study is the first to report a direct causal linkage between the developmental GABA switch and the synaptic plasticity of excitatory transmission in ASD-like pathogenesis. Cytoplasmic chloride ion concentration may be directly associated with synaptic plasticity. Alternatively, chloride cotransporters per se might be functionally linked to the pre- or post-synaptic machinery responsible for LTP. Prenatal exposure to VPA has been reported to impair plastic changes in synaptic transmission in the rat hippocampus (Zhang et al. 2003, Mohammadkhani et al. 2022). On the other hand, conditional knockdown of KCC2 has been reported to impair hippocampal plasticity and memory (Kreis et al. 2023), although the linking machinery has not yet been identified. Dysregulated synaptic plasticity has been assumed to be a critical construct in human ASD pathology (Gilbert and Man 2017), but these arguments are based on molecular associates of ASD-linked mutations, and no direct physiological evidence has been proposed. Molecular cascades responsible for impaired LTP need to be analyzed at the subcellular level, namely the release mechanisms in pre-synaptic terminals and the post-synaptic densities in dendritic spines.

However, several questions remain unanswered. Most importantly, the causal events linking embryonic exposure and post-hatch neuronal malfunction remain unspecified. It also remains unclear which of these neuronal events leads to memory formation impairment. We also have to investigate whether the present findings in chicks are relevant to the human ASD. The following section addresses several of these issues.

### What do VPA and IMI act on the embryonic brain at E14?

As an ASD risk factor, VPA is assumed to act as an inhibitor of histone deacetylases (HDAC; Phiel et al. 2001, Mabunga et al. 2015), as confirmed also in embryonic pallial neurons of chicks (Matsushima et al. 2022). We must be careful about the validity of this assumption because VPA could have a wide spectrum of actions depending on the developmental stage. VPA acts as an anti-epileptic drug through the acute potentiation of GABA actions in the adult central nervous system (Winterer 2003), so that VPA might strongly excite embryonic neurons due to the depolarizing GABA. Similarly, IMI may act as a potent agonist of the nAChR, yet it has a wider spectrum of actions, including altered neurogenesis, migration, and neuroinflammation (Costas-Ferreira and Faro 2021). The simple idea of nonselective excitation of the embryonic neurons is questioned because our previous ballistographic recordings of eggshell vibrations revealed that both VPA and IMI suppressed spontaneous embryonic movements (Matsushima et al. 2022). The responses of embryonic neurons need to be analyzed at the cellular level. For the system level analysis, noninvasive Mn-enhanced MRI imaging will be a powerful tool (Lorenzi et al. 2023).

### How is the depolarizing GABA action maintained after VPA and IMI exposures, while expression of the transporter-coding genes did not differ from control?

RNA-seq and qPCR analyses quantified the mRNA levels but did not address the functional amount of the protein molecules. Protein synthesis, molecular dimerization, and/or cytoplasmic trafficking to the target membrane (Agez et al. 2017) may be impaired after embryonic exposure to risky chemicals, as might be suggested by the increase in RN7SL1 expression. Although the physiological roles of the long non-coding RNA in the normal central nervous system are still largely unknown, recent molecular analyses of post-mortem brains of patients with major depression suicides (Zhou et al. 2018) and Alzheimer’s disease (Liu et al. 2024) suggest their possible involvement in neuroinflammation. Alternatively, functions of the transporters, particularly the neuron-specific KCC2 molecules, might be downregulated by impaired phosphorylation (Watanabe et al. 2019); quantitative phosphoproteomics studies need to be performed, together with specification of the responsible cell types and sub-cellular components.

### Is LTP critical for the formation of imprinting memory or the homeostatic control of E-I balance?

Horn (2004) argued that the biochemical, ultrastructural, and neurophysiological events of IMM are associated with filial imprinting, sharing many features with the process of memory formation in the mammalian hippocampus (Hp). It is highly plausible to assume that the observed LTP, despite being much smaller and fluctuating than that found in mammalian Hp, is critical for some aspects of memory formation. However, this does not imply that LTP represents memory traces. Rather, as for the consolidation processes of episodic memory in the Hp, LTP in the chick IMM could be responsible for the dynamic process of storage relocation from the IMM to elsewhere (Horn 1985).

Alternatively, LTP could be responsible for the homeostatic control of the E-I balance in the synaptic network, as suggested in ASD (Nelson and Valakh 2015). In the present study, we found that embryonic exposure to VPA and IMI deteriorated developmental changes in V_rev_ in P1 chicks. We must examine whether exposures cause long lasting impairments in the post-patch life.

### Do human neurodevelopmental disorders have common pathophysiological constructs with domestic chicks, which are phylogenetically remote animals?

In the rat neocortex, prenatal exposure to VPA has been reported to impar post-natal E-I imbalance via elevated NMDA type glutamate receptor (NMDAR) and the associated molecular machineries including calcium/calmodulin kinase II, hence enhanced LTP (Rinaldhi et al. 2007). In concert, selective blockade of NMDAR acutely rescued some the VPA-induced ASD-like symptoms (memantine acutely injected to 8-16 weeks old mice, Kang and Kim 2015; MK801 injected to post-natal 6-10 days old rats, Mohammadi et al. 2020). The present study in chicks revealed different outcomes of the VPA exposure, namely impaired LTP (**Fig. 3**). Our RNA-seq data also failed to detect significantly different expression of NMDAR subunits (GRIN1, 2A, 2B, 2C, 3A, and 3B; data not shown). NMDARs are actually involved in the imprinting cascade (Horn 2004), and LTP in IMM depends on NMDAR activation (Bradley et al. 1991, Yanagihara et al. 1998). Bath-applied T_3_ (triiodothyronine, a potent activator of imprinting memory formation, Yamaguchi et al. 2012) is shown to suppress NMDAR-mediated plateau potentials in the chick IMM (Saheki et al. 2022). VPA could therefore cause E-I imbalance via different developmental landscapes in taxonomically distant animals such as mammals and birds. Different mechanisms could also underlie LTP, despite the Hebbian nature and critical involvement of NMDAR common in both taxa.

Various nuclear receptors have been suggested as key molecules responsible for neurodevelopmental disorders including ASD (Kong et al. 2020). In particular, the nuclear receptor superfamily 4 group A (NR4A1 and 2) requires attention, as NR4A2 was specified as the chick ortholog of the mammalian cortical layer 5/6 specific genes (Fujita et al. 2019). In situ hybridization revealed high levels of expression in the hyperpallium, mesopallium, and arcopallium of P1 chicks, indicating cell-type homology of the cortical/pallial architectures between birds and mammals. In human cases, haploinsufficiency (Lévy et al. 2018), heterozygous truncation (Song et al. 2022) and de novo mutation (Singh et al. 2020, Jesús et al. 2021) of NR4A2 have been reported to be associated with intellectual disabilities, language impairment and epilepsy. Further brain-wide developmental surveys need be conducted on chicks.

The mTOR signaling cascade is another possible target shared by human ASD (Auerbach et al. 2011) and chick imprinting (Batista et al. 2018). In TSC2 haploinsufficient mouse models, mTOR-dependent spine malformation and the associated increase in spine density have been associated with ASD-like behavioral maldevelopment and cortico-striatal hyperconnectivity (Pagani et al. 2021). In chicks, rapamycin injection acutely blocked imprinting and was accompanied by an increase in mushroom-type dendritic spines in the IMM (Batista et al. 2018). The involvement of NR4A1 nuclear receptors in controlling spine formation after IMI exposure must be considered (Chen et al. 2014, Li et al. 2016). Between mammals and birds, we must accomplish systematic comparisons of the developmental bottleneck, where taxonomic diversification of transcriptional regulators is minimized in the formation of the subcortical visual networks (Matsushima et al. 2024).

## Materials and Methods

### Compliance with ethical standards

The animal experiments were conducted according to the guidelines of the Committee on Animal Experiments of the Health Science University of Hokkaido (approval number #24-042). These guidelines are based on the national regulations for animal welfare in Japan (Law for Humane Treatment and Management of Animals after Partial Amendment No. 68, 2005).

### Animals

Domestic white chickens of a White Leghorn strain (*Gallus gallus domesticus*, locally referred to as “Julia Light”) were used. Fertilized eggs were purchased from Iwamura Point Co. (Yuni, Hokkaido, Japan). Eggs were incubated in the laboratory using type P-008B incubators (Showa Furanki Co., Saitama, Japan) at controlled temperature (37.7 [) and humidity (ca. 80%); inside of the incubator was kept in complete darkness until hatch. All subjects were sexed except those used before hatching.

### Embryonic exposures to risk chemicals

On the embryonic 14 days (E14), eggs received a single injection of 200 μL aqueous solution of chemicals to the air sac chamber through a hole on the round edge of the shell. For experimental treatments, eggs received valproic acid (VPA, 1.16 to 5.82 mg, or 7 to 35 μmole), imidacloprid (IMI, 50 to 250 μg, or 0.2 to 1.0 μmole), or D-tubocurarine chloride pentahydrate (tubocurarine, 0.2 mg, or 0.26 μmole) per egg weighing ca. 50 g. The dosages adopted in the present study were reported to be effective in behavioral analyses (Matsushima et al. 2022). Chemicals were purchased from FUJIFILM Wako Pure Chemical Co. Control eggs received a 200 μL distilled water or no injection.

### Slice preparation

The chicks were housed individually in a dark incubator without light exposure until the experiment. Under anesthesia by intramuscular injection of 0.4 mL of a 1:1 mixture of ketamine (10 mg/mL, Daiichi-Sankyo Pharma Co., Tokyo, Japan) and xylazine (2 mg/mL, Sigma-Aldrich Co., St Louis, USA), the left hemisphere was dissected and immersed in ice-chilled Krebs solution composed of (in mM); NaCl 113.0, NaHCO_3_ 25.0, KCl 4.7, KH_2_PO_4_ 1.2, CaCl_2_ 2.5, MgSO_4_ 1.2, glucose 11.1; pH was adjusted to 7.2-7.4 by bubbling with 95% O_2_ - 5% CO_2_. The left IMM was analyzed because of the functional and morphological asymmetry involved in imprinting memory formation (Horn 1985); specifically, successful training significantly increased the length of the post-synaptic density in spine synapses only in the left IMM (Horn et al. 1985). By using a vibrating micro-slicer (type ZERO-1, Dosaka EM Co., Kyoto, Japan), four 500 μm thick consecutive slices were cut on pseudo-frontal plane roughly corresponding to A9.8 to A8.0 of the chick brain atlas by Kuenzel and Masson (1988) (Fig. 4A). Slices were stored in an interface-type recovery chamber at room temperature (26-27 [) for 1-6 hours. For recording, the slice was placed in a submersion-type chamber (volume ca. 1 mL) that was continuously perfused with Krebs solution held at 29-31 [ at a flow rate of 1.5-2.0 mL per min. The following drugs were applied to the perfusing solution: 6,7-dinitroquinoxaline-2,3-dion (DNQX, 10 μM, Tocris Co.), 1(S),9(R)-(-)-bicuculline methiodide (bicuculline, 10-20 μM, Sigma-Aldrich Co.), VU0463271 (1-10μM, Tocris Co.), bumetanide (20 μg/mL or 55 μM, Tocris Co., in vehicle containing 0.005N NaOH).

### Electrical stimulation and whole-cell intracellular recording

Electrical stimulation was delivered via a concentric metal micro-electrode (O.D. 100 μm, USK-10, Unique Medical Co., Tokyo, Japan) inserted into the slice tissue, with the cathodal center. Constant current (60-200 μA, duration = 100-400 μsec) was supplied by an isolation unit and pulse-generator (SS-203J and SEN-8203, Nihon Koden Co., Tokyo, Japan) at constant intervals (2 or 10 sec). Intracellular whole-cell recordings were performed using nystatin perforation patches. Patch glass pipettes were filled with a solution containing (in mM): K-gluconate 123.0, KCl 18.0, NaCl 9.0, MgCl_2_ 1.0, EGTA 0.2, and HEPES 10.0; the pH was adjusted to 7.3 with KOH. DC-resistance ranged in 5-8 MΩ. Nystatin (Sigma-Aldrich Co., St Louis, USA) was dissolved in DMSO at 50 mg/mL, and 1-5 μL was added to 1 mL of the pipette solution just before backfilling the glass capillaries.

Two types of amplifiers were used. A single-electrode discontinuous voltage clamp amplifier (SEVC, CEZ-3100, Nihon Koden Co., Tokyo, Japan; switching at 20 kHz) and a conventional voltage clamp amplifier (VC, Axopatch 200B, Molecular Devices Co., California, USA). For SEVC recordings, the bridge balance and electrode capacitance compensation were adjusted. For the VC recordings, capacitance and series resistance compensation were not necessary for recording synaptic currents, thus were turned off. The data obtained by these two methods were merged because the similar signals were recorded except for the noise level (VC yielded lower noise than SEVC). The voltage and current records were low-pass filtered at 2 kHz, digitized, and stored using the CED Power 1401 interface and Signal ver.6.02 (digitized at 20 kHz for 25-50 msec per trace; Cambridge Electronic Design Ltd., Cambridge, UK) on a Windows PC. Under microscopic observation, the glass electrodes were inserted at ca. 250 μm ventral to the stimulation electrode (**Fig. 1A**), not necessarily targeting excitatory neurons. The patch recordings were made in a blinded manner, and the recorded neuron types were not identified.

Pilot experiment revealed stable recordings with no detectable changes in GABA reversal potentials up to approximately 30 min after the formation of a giga-ohm seal in control slices, indicating that the present protocol using nystatin perforation did not cause artificial changes in intracellular chloride concentration. A high-chloride pipette solution reversed the GABA current (Sakehi et al. 2022), but this method was not used in this study. In addition, as shown in the vehicle control data in **Fig. 2C** (left), no significant changes were observed in the VPA slices. Data were obtained from excitable neurons (inducible overshooting full action potentials) with initial resting membrane potentials below -30 mV at 3 min after the formation of giga-ohm seal. We discarded the initial records obtained during 10-20 min after the formation of the whole-cell patch. When the input resistance changed by more than 20% of the initial value, we discarded all subsequently obtained data.

### Recording of extracellular field potential

Extracellular field potentials were recorded through a pair of insulated nichrome microwire (O.D. 100 μm). One of the exposed tips (anode) was inserted in the slice tissue, and another (cathode) was submerged in the perfusion solution, and the electrodes were connected to a differential amplifier (MEG-5100, Nihon Koden Co., Tokyo, Japan; gain 0.1 mV/V, band-pass filter 15 Hz - 3 kHz). The cathodal wire was inserted at ca. 250 μm ventral to the stimulation electrode; when accompanied by intracellular recordings, the wire was instead placed at ca. 500 μm ventral (**Fig. 1A, 3A**). As described above, the recorded signals were digitized and stored using the CED Power 1401 and Signal software. Recordings during the initial 10-30 min were discarded until waveform drift ceased. When the amplitude of the pre-synaptic fiber volley changed by more than 10% between the pre-and post-tetanic periods, we discarded all the data obtained from the slice, including the data before the changes.

### RNA-seq and qPCR analysis

Total RNA was extracted from the punched tissue using the FastGene RNA Premium Kit (FG-81050, NIPPON Genetics Co., Tokyo, Japan) and reverse-transcribed to cDNA using the High-Capacity cDNA Reverse Transcription Kit (4368814, Thermo Fisher Scientific Inc., Massachusetts, USA). The expression levels of individual genes were quantified by absolute quantification using KOD SYBR qPCR Mix (QKD-201, TOYOBO Inc., Tokyo, Japan). Standard curves were generated using pGEM-T Easy plasmids containing the amplified target regions. The primers used for standard plasmid construction and qPCR are listed in Supplementary **Table S1**. Gene expression levels were normalized to the expression of glyceraldehyde-3-phosphate dehydrogenase (GAPDH), which served as the internal control.

Total RNA for RNA sequencing was extracted from the punched tissue using a Monarch Total RNA Miniprep Kit (T2010S, New England Biolabs, Inc., Massachusetts, USA), and sequencing was outsourced (NIPPON Genetics Co., Tokyo, Japan). Sequencing reads were trimmed using fastp v0.20.1 and subsequently mapped to the chicken reference genome (GCF_016699485.2) using the STAR v2.3.7a-RSEM v1.3.3 pipeline for expression quantification. Differential expression analyses were performed using the edgeR package in R.

### Statistical analyses

Graphics and statistical computations were performed using RStudio version 3.6.3 (2020-02-29). Conventional parametric tests were conducted. See the Results and Supplementary Materials for details on each data set.

## Acknowledgements

Critical comments and instructive suggestions for the manuscript by Dr. Giorge Vallortigara (University of Trento, CIMeC) are gratefully acknowledged. The valuable instructions on the toxicology of imidacloprid and its metabolites provided by Dr. Yoshinori Ikenaka (Hokkaido University, Faculty of Veterinary Medicine) is also appreciated. We also thank Editage (www.editage.com) for the English language editing.

## Contribution of the authors

T.M. and T.I. conceived the study. T.M. and H.S. performed neurophysiological experiments using slice preparations. N.T. and K.W. designed and performed the molecular biology studies (RNA-Seq and qPCR). T.M. performed statistical analyses. T.M., N.T., K.W., and T.I. wrote the manuscript. All authors agree to publish the paper in its final form.

