## Supplementary materials for "Embryonic exposure to valproic acid and neonicotinoid deteriorates the developmental GABA switch and impairs long-term potentiation in the local circuit of intermediate medial mesopallium of chick telencephalon"

The dataset (raw data in excel format and the RStudio codes in html files) are available at the repository site of the Hokkaido University (HUSCAP)
http://hdl.handle.net/2115/93040

RNA-seq data generated in this study have been deposited to DDBJ Sequence Read Archive (BioProject accession # PRJDB18759) and are publicly available as of the date of publication.

### Contents

Statistical computations
- Fig. 1
- Fig. 2
- Fig. 3
- Fig. 4

Supplementary figures
- Fig. S1
- Fig. S2
- Fig. S3

Supplementary tables
- Table S1
- Table S2
- Table S3

Prepared on September 16, 2024 by Toshiya Matsushima

### Statistical computations

#### Fig. 1 Whole cell recordings of $V_{rev}$ and $V_{rest}$

***Fig.1Ca (Vrev, one-way ANOVA on Vrev: control data, P1, E20 and E18)***
<u>Anova Table (Type II tests)</u>

| Response: Vrev | Sum Sq | Df | F value | Pr(>F) |
|---|---|---|---|---|
| treatment | 2596.6 | 2 | 23.181 | 6.904e-08 *** |
| Residuals | 2856.4 | 51 | | |

***Fig. 1Ca (Vrev, Dunnett multiple comparisons on Vrev: control, VPA, IMI and tubocurarine)***
<u>Fit: aov(formula = Vrev ~ treatment, data = dataset_GABA_reversal_merged_P1)</u>

| Linear Hypotheses: | Estimate | Std. Error | t value | Pr(>\|t\|) |
|---|---|---|---|---|
| 1_VPA – 02_control_P1 == 0 | 8.314 | 1.797 | 4.628 | < 0.001 *** |
| 2_IMI – 02_control_P1 == 0 | 6.594 | 2.164 | 3.047 | 0.00816 ** |
| 3_tubocurarine - 00_control_P1 == 0 | 1.600 | 2.469 | 0.648 | 0.85991 |

***Fig.1Cb (Vrest, one-way ANOVA on Vrest: control data, P1, E20 and E18)***
<u>Anova Table (Type II tests)</u>

| Response: Vrest | Sum Sq | Df | F value | Pr(>F) |
|---|---|---|---|---|
| Treatment | 257.84 | 2 | 3.2731 | 0.04599 * |
| Residuals | 2008.76 | 51 | | |

***Fig. 1Cb (Dunnett multiple comparisons on Vrest: control, VPA, IMI and tubocurarine)***
<u>Fit: aov(formula = Vrest ~ treatment, data = dataset_GABA_reversal_merged_P1)</u>

| Linear Hypotheses: | Estimate | Std. Error | t value | Pr(>\|t\|) |
|---|---|---|---|---|
| 1_VPA - 02_control_P1 == 0 | 2.133 | 1.390 | 1.535 | 0.2989 |
| 2_IMI - 02_control_P1 == 0 | 1.162 | 1.674 | 0.694 | 0.8344 |
| 3_tubocurarine - 00_control_P1 == 0 | 4.200 | 1.910 | 2.199 | 0.0788 |

#### Fig. 2 Whole cell recordings of bumetanide and VU0463271 effects

***Fig. 2C (left; one-sample t-test on $\Delta$Vrev)***
control + VU0426371 (data: delta_Vrev_VU)
t = 6.4293 df = 6 p-value = 0.0006692
VPA+vehicle (data: delta_Vrev_vehicle)
t = -0.27735 df = 7 p-value = 0.7895
VPA+bumetanide (data: delta_Vrev_bumetanide)
t = -3.9659 df = 9 p-value = 0.003275

***Fig. 2C (right: one-sample t-test on $\Delta$Vrest)***
control + VU0426371 (data: delta_Vrest_VU)
t = -0.52759 df = 6 p-value = 0.6167
VPA+vehicle (data: delta_Vrest_vehicle)
t = -0.74067 df = 7 p-value = 0.483
VPA+bumetanide (data: delta_Vrest_bumetanide)
t = -0.22423 df = 9 p-value = 0.8276

#### Fig. 3 Field potential recordings of LTP and fEPSP/presynaptic volley ratio

***Fig.3D LTP is explained by fEPSP/presynaptic volley ratio and treatments***
<u>Anova Table (Type II tests)</u>

| Response: LTP | Sum Sq | Df | F value | Pr(>F) |
|---|---|---|---|---|
| treatment | 0.07890 | 3 | 1.4954 | 0.2312995 |
| fEPSP | 0.18719 | 1 | 10.6436 | 0.0023378 ** |
| treatment:fEPSP | 0.47743 | 3 | 9.0488 | 0.0001193 *** |
| Residuals | 0.66831 | 38 | | |

***Linear regression with interactions between treatment and fEPSP/presynaptic volley ratio***

Call: lm(formula = LTP ~ treatment * fEPSP, data = dataset_LTP_fEPSP_2)

| Coefficients: | Estimate | Std. Error | t value | Pr(>\|t\|) |
|---|---|---|---|---|
| (Intercept) | 1.70645 | 0.13302 | 12.828 | 2.20e-15 *** |
| treatment1_VPA | -0.58511 | 0.19940 | -2.934 | 0.005641 ** |
| treatment2_IMI | -0.74597 | 0.17426 | -4.281 | 0.000122 *** |
| treatment3_tubocurarine | 0.15785 | 0.22502 | 0.701 | 0.487271 |
| fEPSP | 2.1619 | 0.48941 | -4.417 | 8.03e-05 *** |
| treatment1_VPA:fEPSP | 1.95353 | 0.56788 | 3.440 | 0.001427 ** |
| treatment2_IMI:fEPSP | 2.16731 | 0.54708 | 3.962 | 0.000316 *** |
| treatment3_tubocurarine:fEPSP | 0.04122 | 0.70205 | 0.059 | 0.953484 |

***Fig.3E Multiple comparisons of LTP by Dunnett***

Fit: aov(formula = LTP ~ treatment, data = dataset_LTP_2)

| Linear Hypotheses: | Estimate | Std. Error | t value | Pr(>\|t\|) |
|---|---|---|---|---|
| 1_tubocurarine - 0_control == 0 - | 0.02685 | 0.07130 | -0.377 | 0.965 |
| 2_VPA - 0_control == 0 | -0.12620 | 0.07130 | -1.770 | 0.199 |
| 4_IMI - 0_control == 0 | -0.18862 | 0.07130 | -2.646 | 0.030 * |

***Fig.3E Pairwise comparison of LTP between control vs control+VU0463271***

F test to compare two variances, data: LTP by treatment

F = 12.514 num df = 11 denom df = 10 p-value = 0.0003962

Welch Two Sample t-test, data: LTP by treatment

t = 1.4853 df = 12.894 p-value = 0.1615

***Fig.3E Pairwise comparison of LTP between VPA vs VPA+bumetanide***

F test to compare two variances, data: LTP by treatment

F = 0.43281 num df = 11 denom df = 10 p-value = 0.1857

Welch Two Sample t-test, data: LTP by treatment

t = -2.1297 df = 17.067 p-value = 0.04804

***Fig.3F Multiple comparisons of fEPSP/presynaptic volley ratio by Dunett***

Fit: aov(formula = fEPSP ~ treatment, data = dataset_LTP_fEPSP_2)

| Linear Hypotheses: | Estimate | Std. Error | t value | Pr(>\|t\|) |
|---|---|---|---|---|
| 1_VPA - 0_control == 0 | 0.23799 | 0.05075 | 4.689 | < 0.001 *** |
| 2_IMI - 0_control == 0 | 0.16699 | 0.05323 | 3.137 | 0.00867 ** |
| 3_tubocurarine - 0_control == 0 | 0.09215 | 0.05075 | 1.816 | 0.18501 |

***Fig.3F Pairwise comparison of fEPSP/presynaptic volley ratio between control vs control+VU0463271***

F test to compare two variances, data: fEPSP by treatment

F = 0.32145 num df = 11 denom df = 10 p-value = 0.07601

Welch Two Sample t-test, data: fEPSP by treatment

t = -4.1093 df = 15.535 p-value = 0.0008679

***Fig.3F Pairwise comparison of fEPSP/presynaptic volley ratio between VPA vs VPA+bumetanide***

F test to compare two variances, data: fEPSP by treatment

F = 1.177 num df = 11 denom df = 10 p-value = 0.8048

Welch Two Sample t-test, data: fEPSP by treatment

t = 2.6578 df = 20.998 p-value = 0.01472

### Fig. 4 Gene expressions

***Fig. 4Ba Comparisons of slc12a2/slc12a5 ratio (in log scale) among 3 developmental stages in 3 brain regions***

Anova Table (Type II tests), Response: slc12a25_log

| | Sum Sq | Df | F value | Pr(>F) |
|---|---|---|---|---|
| stage | 1.77577 | 2 | 66.2286 | 8.642e-13 *** |
| region | 0.23846 | 2 | 8.8934 | 0.0007266 *** |
| stage:region | 0.32285 | 4 | 6.0204 | 0.0008141 *** |
| Residuals | 0.48263 | 36 | | |

***Fig. 4Bb Comparisons of slc12a2/slc12a5 ratio (in log scale) between control and VPA in 3 brain regions***

Anova Table (Type II tests), Response: slc12a25_log

| | Sum Sq | Df | F value | Pr(>F) |
|---|---|---|---|---|

| | | | | |
|---|---|---|---|---|
| treatment | 0.07932 | 1 | 6.0557 | 0.01984 * |
| region | 0.00439 | 2 | 0.1677 | 0.84641 |
| treatment:region | 0.05302 | 2 | 2.0239 | 0.14979 |
| Residuals | 0.39295 | 30 | | |

---

***Linear regression with interactions between treatment (VPA) and brain regions***

Call: lm(formula = slc12a25_log ~ treatment * region, data = dataset_PCR3)

| Coefficients: | Estimate | Std. Error | t value | Pr(>\|t\|) |
|---|---|---|---|---|
| (Intercept) | -0.04619 | 0.04672 | -0.989 | 0.331 |
| treatmentVPA | 0.02290 | 0.06608 | 0.347 | 0.731 |
| region2_arco | -0.01943 | 0.06608 | -0.294 | 0.771 |
| region3_tectum | -0.06629 | 0.06608 | -1.003 | 0.324 |
| treatmentVPA:region2_arco | 0.03535 | 0.09345 | 0.378 | 0.708 |
| treatmentVPA:region3_tectum | 0.17759 | 0.09345 | 1.900 | 0.067 . |

---

Signif. codes: 0 ‘***’ 0.001 ‘**’ 0.01 ‘*’ 0.05 ‘.’ 0.1 ‘ ’ 1

### Supplementary figures

**Fig. S1** Vrev (GABA reversal potential) was plotted against Vrest (resting membrane potential) for the developmental changes (**A**; E18, E20 and P1), the effect of VPA of 2 doses (**B**; 7 and 35 µmole per egg), IMI of 2 doses (**C**; 50 and 250 µg per egg), tubocurarine (**D**; tubocurarine 0.2mg per egg). Same set of the control P1 data were duplicated in A-D. As the Vrest values considerably overlapped among groups, linear fitting analyses using *lm()* function was performed on Vrev as shown below.

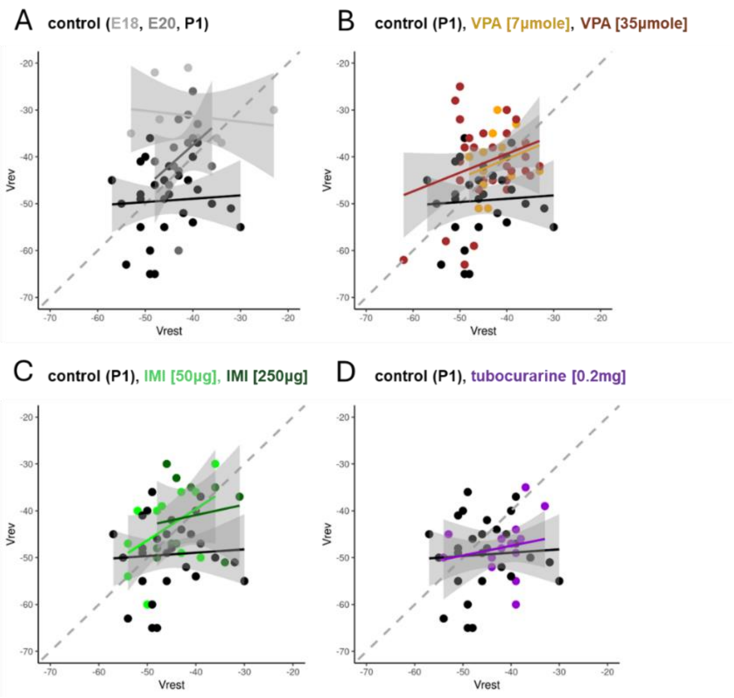


**A: control developmental changes**

Call: lm(formula = Vrev ~ treatment, data = dataset_GABA_reversal_control)

| Coefficients: | Estimate | Std. Error | t value | Pr(>\|t\|) |
|---|---|---|---|---|
| (Intercept) | -49.333 | 1.366 | -36.106 | < 2e-16 *** |
| treatment01_control_E20 | 9.867 | 2.367 | 4.169 | 0.000119 *** |
| treatment02_control_E18 | 18.000 | 2.844 | 6.328 | 6.25e-08 *** |

**B: VPA dose-response**

Call: lm(formula = Vrev ~ treatment, data = dataset_GABA_reversal_VPA)

| Coefficients: | Estimate | Std. Error | t value | Pr(>\|t\|) |
|---|---|---|---|---|
| (Intercept) | -49.333 | 1.467 | -33.621 | < 2e-16 *** |
| treatment1_VPA_7umole | 7.619 | 2.601 | 2.929 | 0.00446 ** |
| treatment2_VPA_35umole | 8.577 | 1.975 | 4.344 | 4.17e-05 *** |

**C: IMI dose-response**

Call: lm(formula = Vrev ~ treatment, data = dataset_GABA_reversal_IMI)

| Coefficients: | Estimate | Std. Error | t value | Pr(>\|t\|) |
|---|---|---|---|---|
| (Intercept) | -49.333 | 1.437 | -34.336 | < 2e-16 *** |
| treatment3_IMI_1ppm | 5.487 | 2.613 | 2.100 | 0.04080 * |
| treatment4_IMI_5ppm | 8.033 | 2.874 | 2.796 | 0.00733 ** |

**D: tubocurarine**

Call: lm(formula = Vrev ~ treatment, data = dataset_GABA_reversal_tubocurarine)

| Coefficients: | Estimate | Std. Error | t value | Pr(>\|t\|) |
|---|---|---|---|---|
| (Intercept) | -49.333 | 1.300 | -37.953 | <2e-16 *** |
| treatment5_tubocurarine | 1.600 | 2.251 | 0.711 | 0.481 |

**Fig. S2** Acute effects of blockers of chloride cotransporters, VU0463271 for KCC2 and bumetanide for NKCC1. Field EPSP responses. Pre-tetanus recording started after the acute effects of the blockers were stabilized, i.e., at 20 min or longer after the onset of infusion. The fEPSP / presynaptic volley ratio recorded in pre-tetanus (x-axis) and the subsequent changes in the fEPSP (26-30 min post-tetanus, y-axis) were plotted. Control and VPA data are reproduced from **Fig. 3D**. Note that larger fEPSP appeared after VU463271, whereas fEPSP was smaller after VPA.

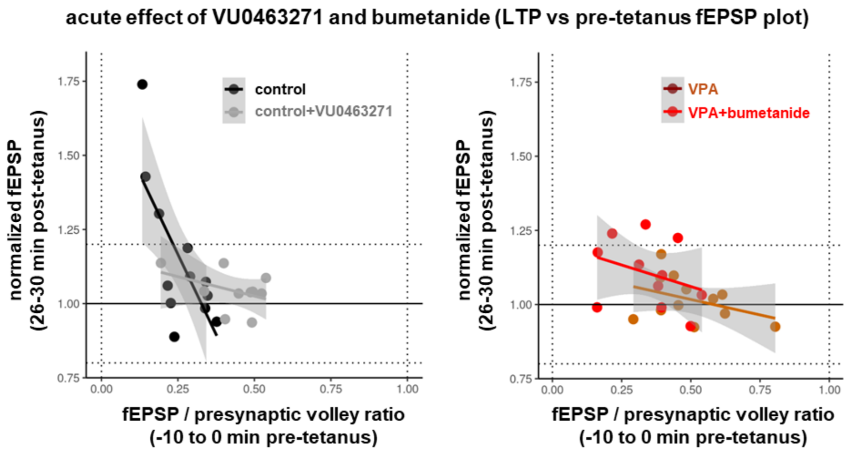

**Fig. S3** qPCR analysis of gene expression profiles. Left raw: nAChR subunits (chhrn4/chrn7 ratio). Middle raw: NeuN gene (rbox3 normalized by gapdh). Right raw: cotransporters NKCC1 by KCC2 (slc12a2/scl12a5 ratio). Log scale. **A)** Developmental changes (stages of E14, E18 and P1) in three regions (IMM, arcopallium and tectum) of untreated samples (5 each). Result of two-way ANOVA are shown in box below each graph, with statistical significance ($p<0.05$) shown in red. Beside the differences among the three regions, significant effects of stages occurred in rbfox3 and slc12a2/scl12a5 ratio. **B)** Comparison between VPA [35µmole/egg] and control P1 chicks (6 each). Two-way ANOVA revealed a significant effect of treatment in slc12a2/scl12a5 ratio, which however did not differ in IMM. **C)** Comparison between tubocurarine [0.2mg/egg] and control P1 chicks (5 each). No significant effect of treatment occurred. Parts of A and B on the slc12a2/slc12a5 ratio (log) are reproduced in Fig. 4B of the main text.

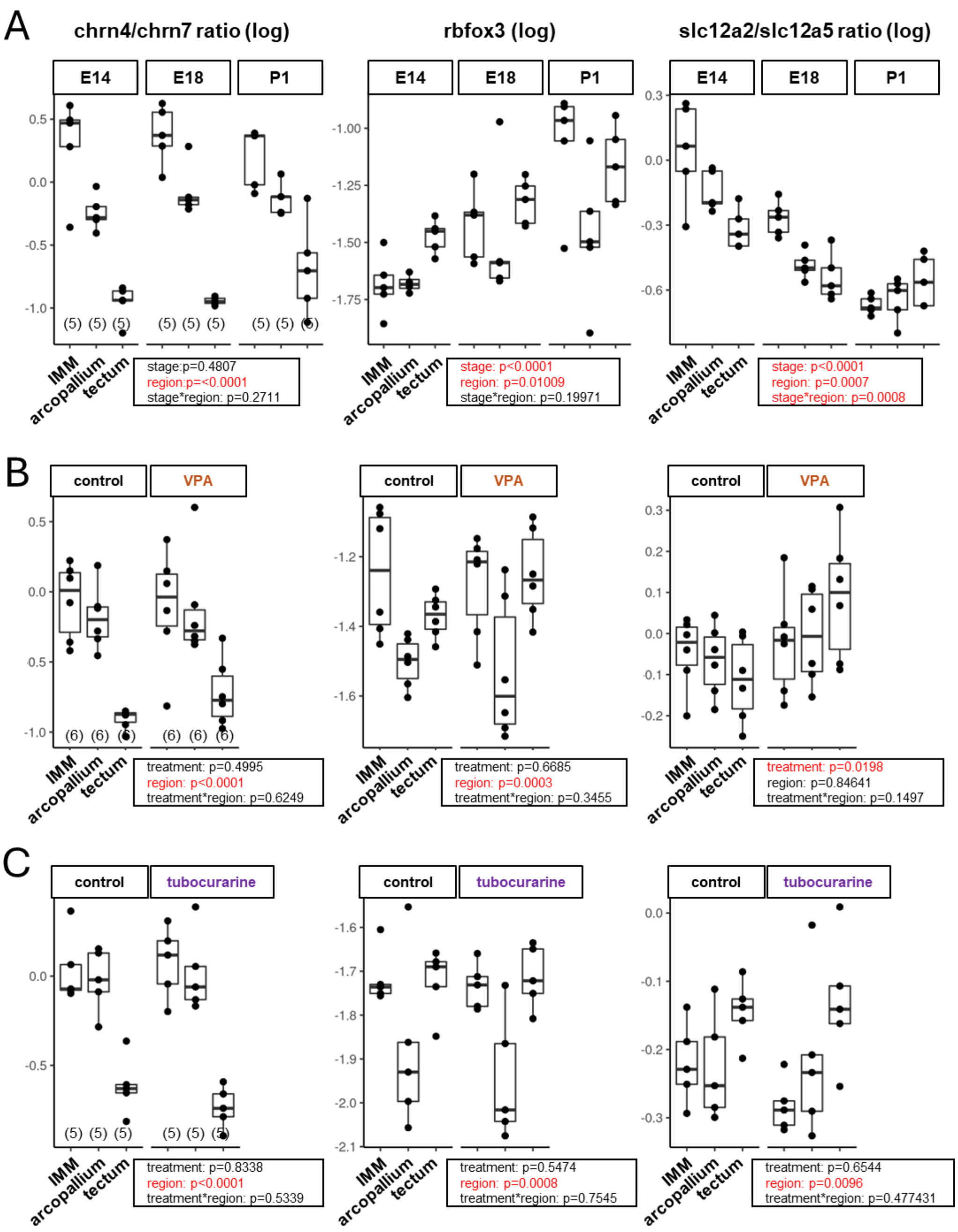

### Supplementary tables

**Table S1**

| gene | accession No. | forward | reverse |
|---|---|---|---|
| chrn4 | NM_204814.1 | TTTCGGATGTGGTCCTGGTC | AAGTCACCATCAGCATTGTTGT |
| chrn7 | NM_204181.2 | TGCAGATCATGGATGTGGATGA | TGGCAGTGTCCCGAAGAATT |
| slc12a2 | XM_040655729.1 | CCAGATGTGAACTGGGGATCA | TTGCCTCCGAGGACCCATAT |
| slc12a5 | XM_040688297.1 | GGTCTTCCTTTGCTGCTCCT | GGGAAGATGTAGGCCAGCAG |
| rbfox3 | XM_040686609.1 | CCGGGTGATGACGAACAAGA | GTACACTCGGCCGTAACTGT |
| gapdh | NM_204305.2 | GCAGATGCAGGTGCTGAGTA | GACACCCATCACAAACATGG |

**Table S2** RNA-seq data of IMM after embryonic exposure to VPA and IMI. Those genes with FDR (false discovery rate) <0.1 were shown. Those genes indicated in red (and blue) were similarly upregulated (and downregulated) in both of the VPA and IMI. Genes are ordered according to the ratio to the control. Genes labelled as LOCxxxx represent uncharacterized genes.

| gene name | NCBI symbol | VPA | IMI |
|---|---|---|---|
| WIPF3-like | LOC107055396 | 7.472968 | 10.09147 |
| LOC124417499 | LOC124417499 | 7.405521 | 9.028126 |
| ERVK-10-like | LOC121107320 | 7.378221 | 7.02123 |
| RN7SL1 | RN7SL1 | 5.366336 | 4.444392 |
| LOC121113304 | LOC121113304 | 2.534067 | 3.29161 |
| SNORA73 | LOC112530237 | 6.792813 | 0 |
| KCNQ1 | KCNQ1 | 6.635323 | 0 |
| LOC124418341 | LOC124418341 | 6.495934 | 0 |
| LOC101748814 | LOC101748814 | 4.639356 | 0 |
| QRFPR | QRFPR | 4.044614 | 0 |
| TMEM233 | TMEM233 | 3.969052 | 0 |
| HBA1 | HBA1 | 1.6945 | 0 |
| LOC121106571 | LOC121106571 | 0 | 6.775907 |
| PHF7-like | LOC112533357 | 0 | 6.584688 |
| CDCA8-like1 | LOC107057183 | 0 | 5.985043 |
| LOC121111083 | LOC121111083 | 0 | 5.970251 |
| LOC121113405 | LOC121113405 | 0 | 5.922122 |
| LOC124417295 | LOC124417295 | 0 | 5.029757 |
| LOC770574 | LOC770574 | 0 | 4.030542 |
| LOC107054402 | LOC107054402 | 0 | 3.466611 |

| | | | |
|---|---|---|---|
| ODF3-like | LOC107050025 | 0 | 2.984196 |
| LOC121109595 | LOC121109595 | 0 | 2.853978 |
| LOC107053944 | LOC107053944 | 0 | 2.674105 |
| LOC107052086 | LOC107052086 | 0 | 2.604105 |
| LOC771161 | LOC771161 | 0 | 2.286273 |
| LOC107049018 | LOC107049018 | 0 | 2.202385 |
| LOC107049870 | LOC107049870 | 0 | 2.169354 |
| AASDH | AASDH | 0 | 1.834962 |
| OLIG2 | OLIG2 | 0 | -1.59196 |
| CEBPD | CEBPD | 0 | -1.60744 |
| RASL11A | RASL11A | 0 | -1.98416 |
| CYR61 | CYR61 | 0 | -2.01496 |
| DOK1 | DOK1 | 0 | -2.07236 |
| NR4A1 | NR4A1 | 0 | -2.19152 |
| FOS | FOS | 0 | -2.26126 |
| DDIT4 | DDIT4 | 0 | -2.30196 |
| IL1B | IL1B | 0 | -2.39659 |
| TAC1 | TAC1 | 0 | -2.55539 |
| THRSPB | THRSPB | 0 | -2.86232 |
| MHCYL | MHCYL | 0 | -4.47313 |
| CLDN1 | CLDN1 | 0 | -6.68059 |
| LOC107054815 | LOC107054815 | 0 | -6.70821 |
| CDCA8 | LOC121107214 | 0 | -8.40193 |
| LOC112532278 | LOC112532278 | 0 | -8.94964 |
| NR4A2 | NR4A2 | -1.74592 | 0 |
| CDCA8-like2 | LOC107053791 | -2.23743 | 0 |
| DDC | DDC | -3.40199 | 0 |
| CAPG | CAPG | -4.85601 | 0 |
| LOC121111356 | LOC121111356 | -6.90277 | 0 |
| CDCA8-like3 | LOC107057197 | -8.35058 | 0 |
| PPP1RL | PPP1RL | -11.2543 | 0 |
| MHCY2B7 | MHCY2B7 | -7.81637 | -5.03253 |
| LOC121106939 | LOC121106939 | -5.41817 | -8.02846 |
| LOC107049675 | LOC107049675 | -8.50422 | -8.48286 |
| LOC124417862 | LOC124417862 | -11.0743 | -11.0533 |

**Table S3** SLC12A2/5 expressions in IMM after embryonic treatment by VPA and IMI. Data obtained from 6 individual chicks (each 4 tissue punchouts) are shown.

| gene | ctrl_1 | ctrl_2 | VPA_1 | VPA_2 | IMI_1 | IMI_2 |
|---|---|---|---|---|---|---|
| NKCC1: SLC12A2 | 4.33 | 7.98 | 7.15 | 6.51 | 8.17 | 6.49 |
| KCC2: SLC12A5 | 89.24 | 51.66 | 44.95 | 49.69 | 49.00 | 32.32 |

Ratio of SLC12A2 by SLC12A5

| ctrl_1 | ctrl_2 | VPA_1 | VPA_2 | IMI_1 | IMI_2 |
|---|---|---|---|---|---|
| 0.0485 | 0.1545 | 0.1591 | 0.1310 | 0.1667 | 0.2008 |

Four isoforms of SLC12A2 and two of SLC12A5.

| | | ctrl_1 | ctrl_2 | VPA_1 | VPA_2 | IMI_1 | IMI_2 |
|---|---|---|---|---|---|---|---|
| NKCC1: SLC12A2 | NKCC1: XM_003643059.6 | 2.47 | 4.68 | 4.14 | 4.03 | 4.69 | 2.53 |
| NKCC1: SLC12A2 | NKCC1: XM_004949377.5 | 0.22 | 0.00 | 0.42 | 0.00 | 0.49 | 0.00 |
| NKCC1: SLC12A2 | NKCC1: XM_004949378.5 | 1.65 | 2.76 | 2.59 | 2.19 | 2.99 | 3.96 |
| NKCC1: SLC12A2 | NKCC1: XM_040655729.2 | 0.00 | 0.54 | 0.00 | 0.29 | 0.00 | 0.00 |
| KCC2: SLC12A5 | KCC2a: XM_025142286.3 | 3.92 | 5.18 | 0.88 | 1.00 | 1.49 | 0.01 |
| KCC2: SLC12A5 | KCC2b: XM_025142287.3 | 85.32 | 46.48 | 44.07 | 48.69 | 47.52 | 32.31 |

Ratio of SLC12A2 isoform (XM_004949378.5) by SLC12A5 isoform (XM_025142287.3)

| ctrl_1 | ctrl_2 | VPA_1 | VPA_2 | IMI_1 | IMI_2 |
|---|---|---|---|---|---|
| 0.0193 | 0.0594 | 0.0588 | 0.0450 | 0.0629 | 0.1226 |
